# Tumor suppressor NME1/NM23-H1 modulates DNA binding of NF-κB RelA

**DOI:** 10.1101/2024.10.06.616908

**Authors:** Shandy Shahabi, Mano Maurya, Shankar Subramaniam, Gourisankar Ghosh

## Abstract

The dimeric NF-κB family of transcription factors activates transcription by binding sequence-specifically to DNA response elements known as κB sites, located within the promoters and enhancers of their target genes. While most NF-κB remain inactive in the cytoplasm of unstimulated cells, a small amount of RelA, one of its members, persists in the nucleus, ensuring low-level expression of genes essential for homeostasis. Several cofactors have been identified that aid in DNA binding of RelA. In this study, we identify NME1 (nucleoside diphosphate kinase 1) as a cofactor that enhances RelA’s ability to bind κB sites within the promoters of a subset of its target genes, promoting their expression under both unstimulated and stimulated conditions. Depletion of NME1 influences activation or repression of several genes that are unresponsive to TNFα, despite containing κB sites in their promoters but not in clusters. This suggests that clustering of kB sites may be necessary for RelA-dependent transcription complex assembly. NME1 appears to act as a cofactor for other transcription factors to regulate these genes. NME1 does not directly contact κB DNA but interacts with RelA, with this interaction being further strengthened in the presence of κB DNA. Notably, NME1 alone has a marginal effect in enhancing RelA’s DNA binding, suggesting that NME1 likely cooperate with other cofactors to regulate DNA binding and transcription through RelA. These observations underscore the intricate assembly of transcription complexes centered on NF-κB.

## INTRODUCTION

DNA-binding transcription factors (TFs) activate or repress transcription by binding to specific sequences within promoters/enhancers of target genes. DNA binding of these TFs has been studied *in vitro* using primarily purified DNA binding domains (Lambert et al., 2018; Todeschini et al., 2014). However, full-length TFs often exhibit poor DNA binding affinity *in vitro*, complicating the understanding of their precise DNA binding mechanism, especially when their nuclear concentrations are low. Additionally, *in vitro* DNA binding experiments are often being done under non-physiological conditions, which also do not reflect the binding in cells. However, a plethora of studies have demonstrated that conclusions made from *in vitro* experiments mostly capture *in vivo* functions. In this study, we use NF-κB transcription factors as a model to explore these outstanding issues related to DNA binding by TFs *in vitro* and in cells.

The NF-κB family, crucial for immune responses, inflammation, and cancer, includes five protomers: p50/NF-κB1, p52/NF-κB2, RelA/p65, cRel, and RelB (Ghosh and Hayden, 2012; Wang et al., 2012). These form various dimers that bind 10-bp κB sites with the consensus 5’-GGGRN<u>N</u>YYCC-3’. Despite the consensus, many κB sites deviate significantly, indicating weak binding might still suffice for transcriptional activation *in vivo* (Siggers et al., 2011). Numerous studies suggest that NF-κB relies on other supporting factors (cofactors) to enhance its transcriptional activity, even enhancing DNA binding activity in some cases (Mulero et al., 2019). Cofactors, often non-DNA binding proteins, interact with NF-κB away from the DNA binding region. An early identified cofactor is IRF3, which interacts with the RelA homodimer to activate the expression of Cxcl-10 in response to LPS (Leung et al., 2004). Although IRF3 is a legitimate DNA binding transcription factor, in this case, it acts remotely from the DNA surface and does not require its DNA binding domain. What is remarkable is that IRF3 requires the presence of an A:T bp at a specific position instead of T:A to bind RelA (Leung et al., 2004). Other documented cofactors include RPS3, Sam68, and nucleoplasmin/NPM1 (Fu et al., 2013; Lin et al., 2017; Wan et al., 2007). Although the κB sites and corresponding genes controlled by these cofactors have been identified, the features in the κB sites necessary for a cofactor have not been properly examined. Two of the better characterized cofactors are RPS3 and Sam68. RPS3 activates a subset of genes such as IL-8 and IκBα in T cells in response to PMA but does not activate CD25. On the other hand, Sam68 activates CD25 but not IL-8 (Wan et al., 2007). In all these cases, cofactors stabilize the NF-κB:DNA complex (Fu et al., 2013). Nothing is known about the mechanism by which a cofactor stabilizes specific complexes between a κB site and an NF-κB dimer. We showed that the DNA binding Rel homology region of NF-κB dimers exhibits robust DNA binding under non-physiological low ionic strength but loses its affinity with increasing ionic strength. Surprisingly, at physiological ionic strength, full-length NF-κB dimers bind κB DNA strongly, and affinity increases in the presence of RPS3 by directly interacting with both the RHR and transcription activation domain (TAD) (Mulero et al., 2018). We have recently reported that a host of protein factors associate with RelA on promoters. Among them are the transcription factors which play a stabilizing role in recruiting RelA at the transcription sites with help from a large number of κB sites. Many of these sites display weak affinity for RelA *in vitro*.

There are also cofactors that destabilize NF-κB:DNA complexes and repress gene transcription. One such repressing or negative cofactor has been reported that represses PD-L1 expression by NF-κB RelA. This cofactor is retinoblastoma (Rb) protein, which, upon phosphorylation by CDK4, interacts with RelA at a loop further away from the DNA binding surface and removes RelA from the DNA (Jin et al., 2019). However, in none of these cases is the mechanism of how *in vivo* specificity is achieved through selective recognition of certain κB sites while discriminating against others known.

NF-κB RelA dimers also maintain the expression of several genes under unstimulated conditions, including the Nfkbia and Nfkbie genes, which code for the IκBα and IκBε inhibitors of NF-κB (Hoffmann et al., 2002). It was previously reported that low levels of RelA are present in the nuclei of resting cells, which are responsible for the homeostatic expression of these genes (Mathes et al., 2008; O’Dea et al., 2007). This is particularly intriguing as to how RelA accomplishes DNA binding under these conditions. We reasoned that different sets of cofactors must be present in unstimulated cells to support RelA’s DNA binding since RelA’s nuclear concentration is significantly low in unstimulated cells. We set out to identify new cofactors of RelA using biochemical methods and determine the genes that are regulated by a new pair of RelA and cofactor. We identified nucleoside diphosphate kinase 1 encoded by the gene NME1 (also known as NM23-H1) as a cofactor of RelA in unstimulated cells. We found that only a subset of genes is affected by NME1 binding. The κB sites of these genes show a common feature of partial NF-κB consensus. We found that the effect of NME1 in recruiting RelA to DNA is marginal when it acts alone; however, in the presence of other unknown cofactors, NME1 is effective. It is as if there are tens, if not hundreds, of cofactors assisting in NF-κB recruitment.

## RESULTS

### Isolation of RelA-specific transcriptional cofactors in unstimulated cells

To confirm the presence of RelA-specific cofactors in the nuclear extracts of unstimulated HeLa cells, we performed EMSA using four different but well known κB sites (MHC-κB, IFNβ-κB, PSel-κB, and Ig-κB). Each of these κB sites is known to have some preference for different NF-κB dimers (Ghosh et al., 2012; Mulero et al., 2019; Pan et al., 2023; Siggers *et al*., 2011). The indicated amounts of recombinant RelA homodimer and p50:RelA heterodimer showed very little shifted NF-κB:DNA complex (**Figure 1A** and **1B**). However, the addition of 100 ng of unstimulated nuclear extract significantly enhanced the binding of these dimers to DNA. The presence of the RelA subunit in the shifted complex was confirmed by a supershift with an anti-RelA antibody (**Figure 1A**). These results indicate the presence of factors in the nuclear extract that promote RelA’s ability to bind DNA. Interestingly, the NF-κB:DNA complex did not shift further in the gel compared to the pure RelA complex, suggesting that the enhancer(s) in the nuclear extract interact transiently with NF-κB or both NF-κB and DNA in the NF-κB:DNA binary complex.

**Figure 1.**
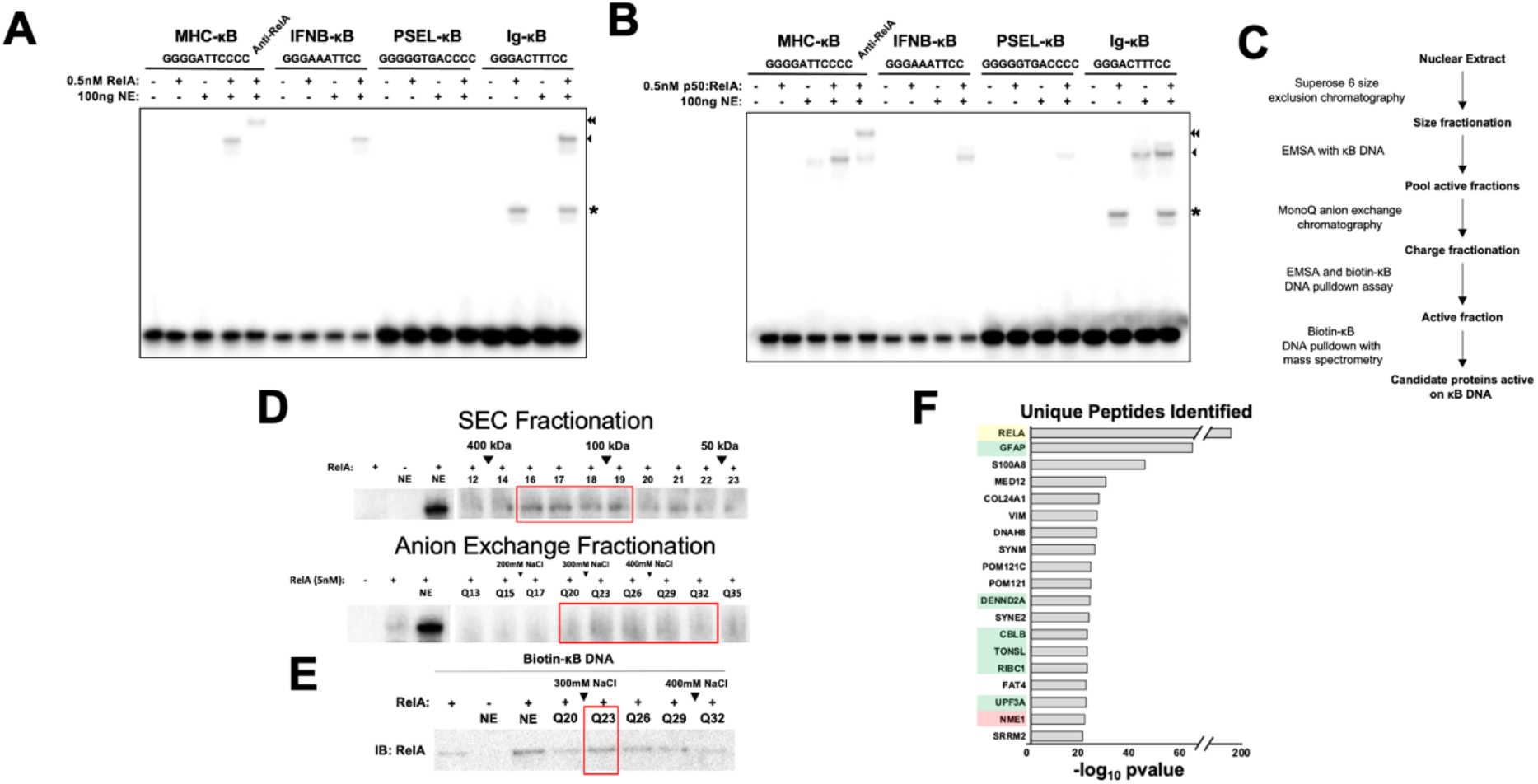
Isolation of RelA-specific transcriptional cofactors in unstimulated cells. **A**. and **B**. EMSA assay with recombinant RelA homodimer (**A**.) or p50:RelA heterodimer (**B**.) and unstimulated HeLa nuclear extract. DNA sequences used in reaction are indicated above (Asterisk is non-specific binding from nuclear extract, single arrow is binding of RelA:RelA or p50:RelA to κB DNA, and double arrow is supershifted band with RelA-specific antibody). **C**. Unstimulated HeLa nuclear extract from 3 15cm plates was collected and first fractioned by size-exclusion chromatography. Fractions were assayed by EMSA and active fractions were pooled and further fractionated by anion exchange chromatography. A biotin-κB DNA streptavidin bead pulldown was then performed on the active fraction and analyzed by mass-spec. **D**. EMSA binding analysis of recombinant FL-RelA with nuclear extract fractioned by size-exclusion (top) and anion exchange (bottom). **E**. Active anion exchange fractions were assayed by biotinylated DNA-streptavidin agarose pulldown with recombinant FL-RelA, and precipitated RelA was analyzed by Western blot with RelA antibody. **F**. Barplot representation of pvalue for peptides identified by mass-spec in RelA-dependent biotin-κB DNA pulldown with streptavidin beads. Proteins listed were not identified in control pulldown without RelA (Yellow is RelA used as bait, green are proteins targeted for subsequent knockdown, and red is NME1 that is further characterized).

We further investigated whether nuclear factors facilitate the DNA binding of other NF-κB dimers by testing the p50 and p52 homodimers on the same κB sites (**Supplementary Figure 1**). Nuclear extracts only marginally enhanced the complex formation when the p50 homodimer was bound to the MHC-κB or PSel-κB, but no enhancement was observed for Ig-κB and IFNb-κB sites. In contrast, the p52 homodimer exhibited distinct behavior: Its binding to MHC-κB and PSel-κB sites resulted in slower mobility complexes, we previously reported that one of these slower mobility complexes contained Bcl3 (Wang et al., 2012). However, p52’s binding to Ig-κB and IFNβ-κB sites was weak, and nuclear extracts did not further enhance the complex formation. These results suggest that DNA binding of RelA as homo- and heterodimer with p50 is more strongly influenced by factors in nuclear extracts compared to the p50 and p52 homodimers.

To identify novel RelA-specific cofactors in the nucleus, we attempted to purify them using three chromatographic steps: size exclusion, anion-exchange and affinity. Following each step of purification, fractions were assessed for their ability to support RelA’s binding to DNA by EMSA. For the affinity chromatography, we employed a biotinylated κB DNA sequence derived from the promoter of interferon beta (IFNβ-κB). Notably, we observed a progressive loss in the compactness of the shifted RelA complex after each purification step, indicating that multiple factors contribute of the RelA interaction with DNA and some of them are being lost during purification.

The purification scheme and the corresponding activity of fractions are shown in **Figures 1C** and **1D**. After the second step, the active fractions were pooled and subjected to affinity chromatography using a biotinylated IFNβ-κB DNA in the presence of RelA homodimer. As a control, we performed the pulldown experiments in the absence of RelA homodimer. We verified using WB that more RelA remained bound to the biotinylated κB DNA in the presence of fraction Q23 (**Figure 1E**). Proteins eluted from the κB DNA affinity beads with and without RelA were then subjected to MS-MS analysis. In total, sixteen proteins were identified in the IFNβ-κB bound fractions that were not present when RelA homodimer was omitted in the pulldown reaction (**Figure 1F**). Among these, nine were abundant cellular proteins likely binding non-specifically to κB DNA and were not considered for further study.

### NME1 enhances DNA binding of RelA

We further investigated the role of seven remaining proteins (GFAP, DENND2A, CBLB, TONSL, RBC1, UPF3A and NME1). We performed knockdowns (KD) of the corresponding genes and assessed their impact on RelA’s DNA binding. KD efficiency for NME1 was greater than 75% as judged by mRNA and protein levels (**Supplementary Figure 2A**). Unstimulated nuclear extracts from each knockdown cell line were prepared, and the ability of these extracts to enhance RelA binding was tested. While most knockdowns had marginal effects, we consistently observed that knockdown of NME1 (also known as NM23-H1) and to some extent DENND2 led to reduced binding of RelA to DNA, indicating a potential role for these proteins in stabilizing the RelA:DNA complex (**Supplementary Figure 2C**). NME1 was discovered as an anti-metastatic protein (hence the other name non-metastatic protein 23 H1; NM23-H1) (Puts et al., 2018). Subsequently, many of its other functions such as exonuclease and histidine kinase activities were identified. A close homolog of NME1 (NME2/NM23-H2) was shown to regulate transcription by binding to G-quadruplexes (Sharma et al., 2021).

Importantly, NME1 KD had minimal impact on either IκBα degradation or nuclear RelA levels (**Supplementary Figure 2C-D**). However, DENND2 KD affected IκBα levels suggesting its complex role in the NF-κB regulation pathway (not shown). We decided not to continue with DENND2A any further. We proceeded to investigate the precise role of NME1 in supporting the RelA:DNA complex formation in NME1 KD HeLa cells. We treated NME1 or control (scr) KD HeLa cells with TNFα for 0, 15, 30 and 60 min and generated nuclear extracts from these cells. We reproducibly found that nuclear extracts made with NME1 KD HeLa cells at early time of induction (15 and 30 min) partly failed to support RelA’s DNA binding but not the extract generated at 60 min of TNFα induction (**Figure 2A**). We also tested the effect of increasing amounts of unstimulated HeLa nuclear extracts isolated from scr and NME1 KD cells on κB DNA binding by recombinant RelA homodimer (**Figure 2B**). The nuclear extract isolated from scr KD cells showed slightly larger enhancement of RelA’s DNA binding compared to NE from NME1 KD cells. Additionally, we assessed the DNA binding ability of transfected HA-RelA expressed in scr and NME1 KD HeLa cells. A significant reduction in DNA binding was observed when NME1 was depleted, despite slightly higher nuclear RelA levels in these cells (**Figure 2C**). We next tested if NME1 overexpression could induce reporter gene expression driven by κB sites. The luciferase reporter assay confirmed that overexpressed NME1 induced the reporter expression (**Figure 2D**).

**Figure 2.**
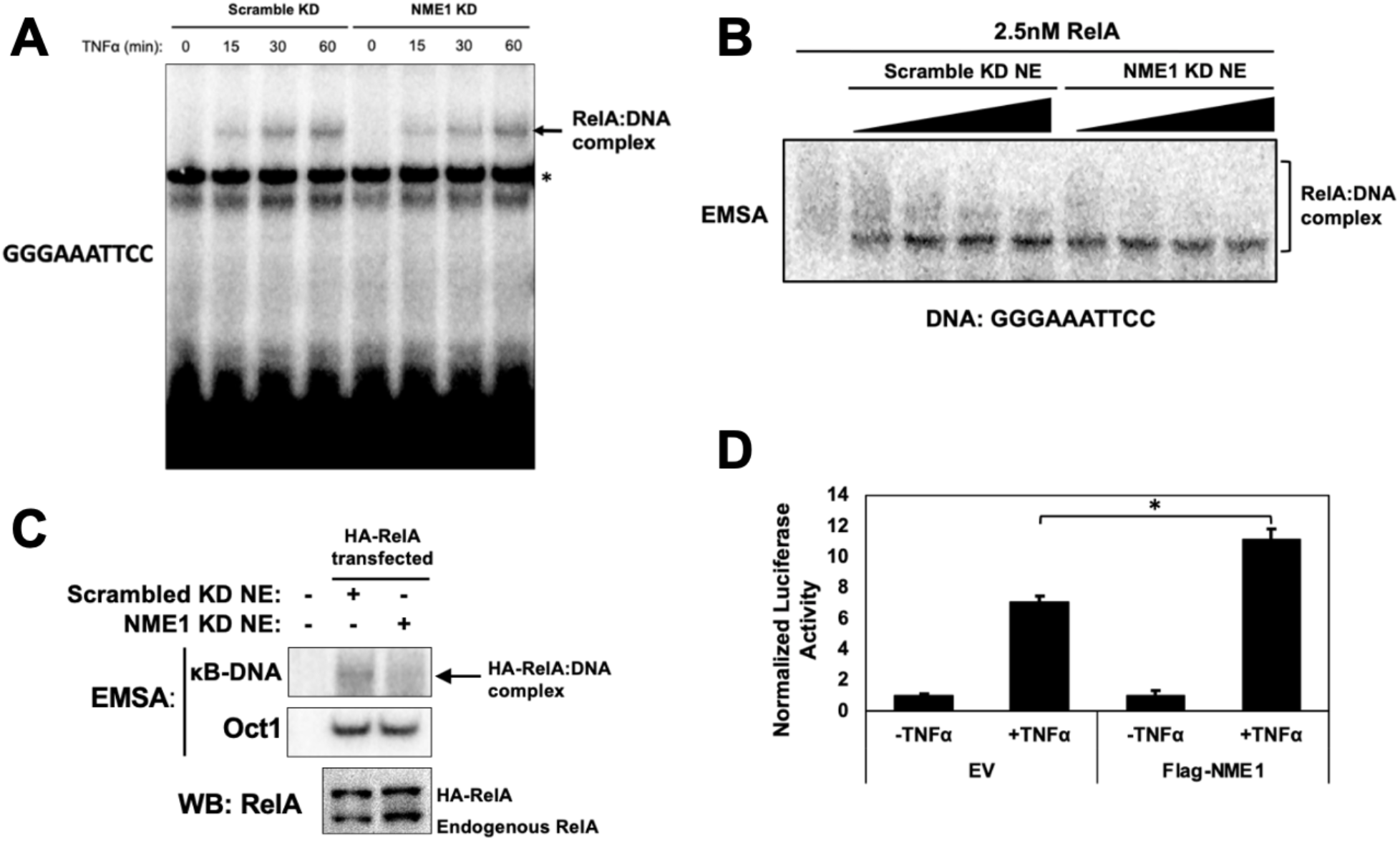
Knockdown of NME1 reduces RelA DNA binding. **A**. EMSA analysis of endogenous NF-κB binding in nuclear extract from stable scramble or NME1 knockdown HeLa cell lines. Cells were stimulated with 20 ng/mL TNFα for the indicated timepoints prior to preparation of nuclear extract. **B**. EMSA with recombinant FL-RelA and increasing amounts of unstimulated nuclear extract from scramble or NME1 knockdown HeLa cell lines. **C**. EMSA analysis of nuclear extract from stable scramble or NME1 knockdown HeLa cell lines transfected with HA-tagged RelA. **D**. Luciferase activation assay with a NF-κB driven luciferase construct cotransfected with empty vector or FLAG-tagged NME1 in HEK293T. Cells were stimulated for 8 hours with control DMSO or TNFα prior to preparation of lysate. AT16 κB DNA was used luciferase construct. Luciferase reading were normalized to Renilla internal control and values are represented as mean ± SD of three independent experimental replicates.

To better understand the mechanism by which NME1 enhances RelA’s DNA binding, we tested whether NME1 and RelA interact (**Figure 3A**). Immunoprecipitation experiments in cotransfected HEK293T cells confirmed an interaction between these two proteins. We further prepared recombinant NME1 from *E. coli* expression system and tested the effect of pure NME1 on interaction with RelA in the presence and absence of κB DNA (**Figure 3B**). EMSA revealed that recombinant NME1 enhanced DNA binding by recombinant RelA *in vitro* (**Figure 3C**). Additionally, a GST pulldown experiment using recombinant GST-NME1 and His-RelA demonstrated a weak yet direct interaction between these two proteins (**Figure 3D**). Interestingly, the presence of IFNβ-κB DNA enhanced this interaction, while mutant DNA did not (**Figure 3E**). Taken together, these results confirm that NME1 stabilizes the RelA:DNA complex and augments transcriptional activity of RelA by making direct protein-protein contacts. However, the interactions between NME1 and RelA appeared to be relatively weak and the stabilization effect on the RelA:DNA complex was marginal.

**Figure 3.**
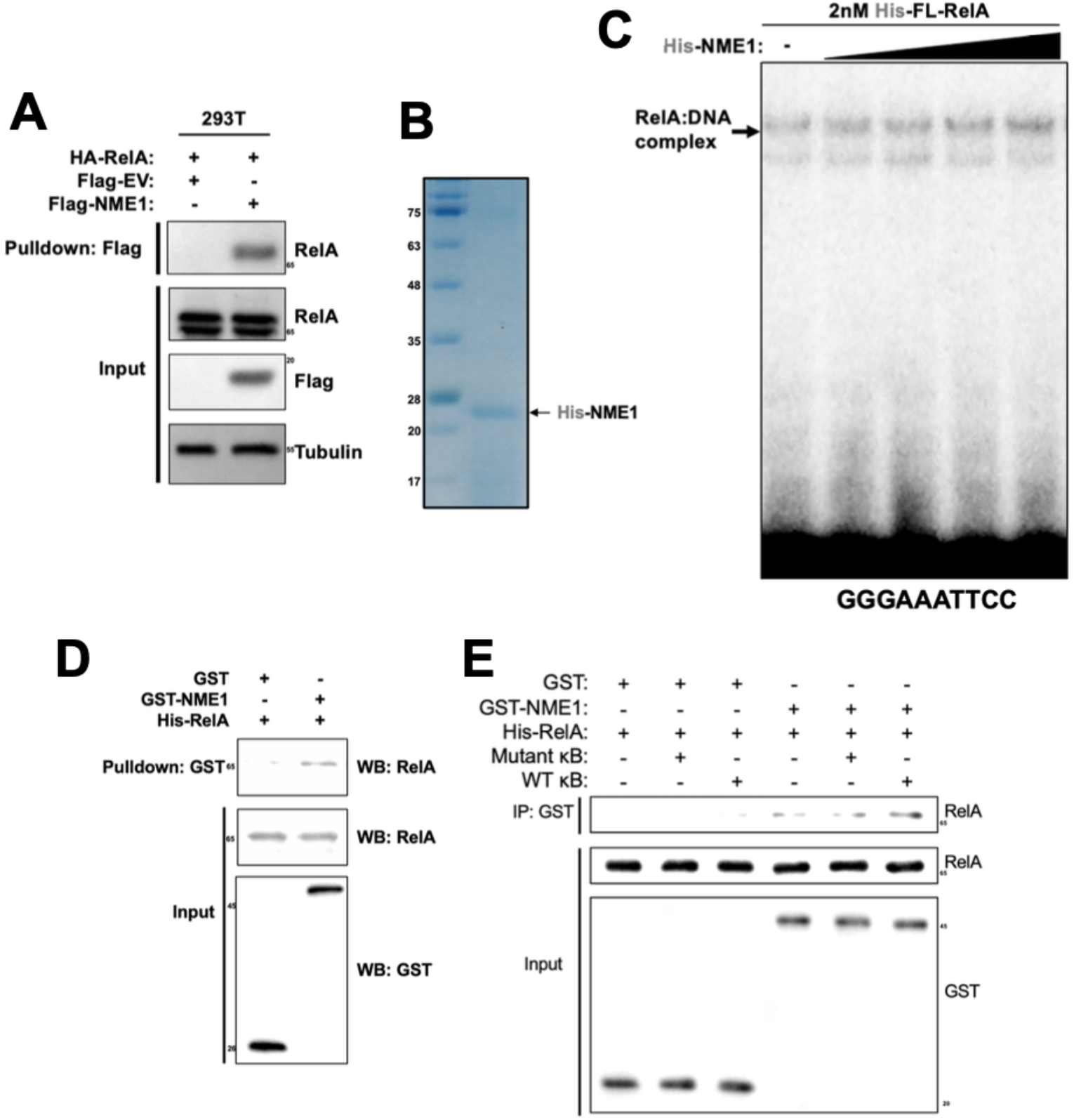
NME1 interacts directly with RelA and enhances binding affinity *in vitro*. **A**. FLAG pulldown assay in whole cell lysate from empty vector or FLAG-NME1 transfected HEK293T. Cells were cotransfected with HA-tagged RelA and precipitated proteins were analyzed by Western blot using the indicated antibodies. **B**.Coomassie stained protein gel of purified His-tagged NME1 from *E. coli*. **C**.EMSA assay with recombinant FL-RelA and increasing amounts of recombinant His-NME1. The sequence of the radiolabeled κB DNA probe is indicated below. **D**. GST-pulldown assay with purified GST or GST-NME and FL-RelA. **E**. GST-pulldown assay with GST or GST-NME1 and FL-RelA in the presence or absence mutated κB DNA or wild type κB DNA

### Effect of NME1 on promoter-specific DNA binding and gene expression in TNFα-stimulated cells

To understand how NME1 affects gene regulation by RelA, we performed RNA sequencing experiments (RNA-seq) in unstimulated and TNFα-stimulated HeLa cells with either scr or NME1 KD. RNA was isolated from each of the four experimental conditions in duplicate, followed by library preparation, sequencing, and data analysis. The heatmap and volcano plot present genes that are either upregulated or downregulated upon NME1 KD compared to scr control (**Figure 4A and 4B**). Pathway analysis revealed a significant upregulation of genes involved in cytokine and inflammatory response processes upon TNFα stimulation, as expected (**Figure 4C**). Not surprisingly, the majority of these genes belong to cytokine, chemokine, and IκB families with κB sites located near the transcription start sites. Gene ontology analysis of up or down regulated genes upon NME KD in unstimulated cells revealed enrichment of genes involved in various biological processes, notably mitosis and cell division in genes that are downregulated and nucleotide and metabolic pathways in upregulated genes (**Supplementary Figures 3A and 3B**).

**Figure 4.**
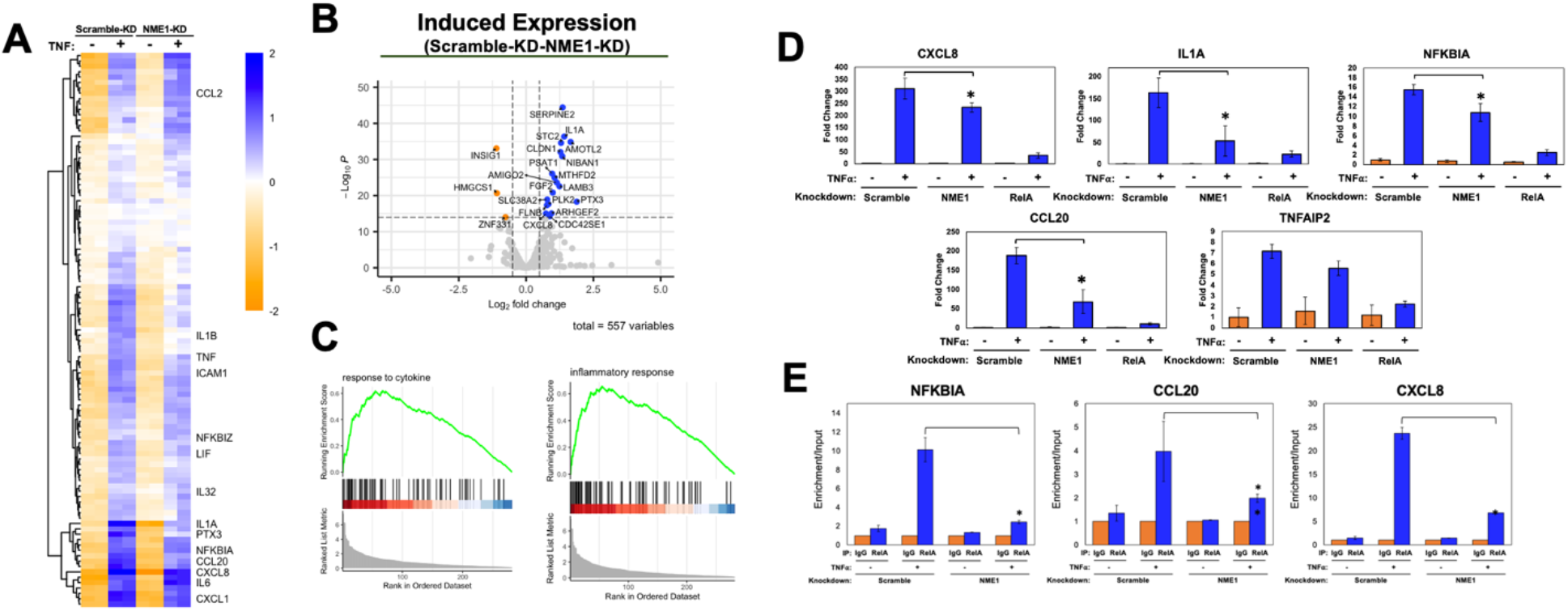
NME1 modulates DNA binding and transcriptional regulation of RelA upon TNFα stimulation. **A**. Heatmap representation of unsupervised hierarchical clustering of differentially expressed TNFα inducible genes (n = 103;) from RNA-seq of scramble and NME1 knockdown HeLa cells stimulated for 1 hour. Genes were identified as TNFα dependent if log_2_ fold change > 1 and p-value < 0.005 in scramble control cells following stimulation. Color represents distance of log transformed transcript counts from mean. **B**. Volcano plot representing differentially expressed TNFα inducible genes between scramble and NME knockdown in stimulated cells. **C**. Gene set enrichment analysis plots of top pathways identified in TNF stimulated control cells compared to unstimulated cells. **D**. RT-qPCR analysis of differentially expressed genes in stable scramble, NME1, and RelA knockdown HeLa cell lines following 1 hour TNFα stimulation. Expression levels were normalized to GAPDH and values are represented as mean ± SD of three independent experimental replicates; * = p value < 0.05. **D**. RelA ChIP-qPCR in scramble and NME1 knockdown HeLa cells stimulated with TNFα for 30 minutes. Enrichment was normalized to input and values are represented relative IgG in three replicates’ * = p value < 0.05.

We observed a reduction in a subset of known RelA target gene expression upon NME1 KD following TNFα stimulation. To validate these observations, HeLa cells with scr KD, NME KD, and RelA KD were treated with TNFα or control DMSO for 1 hr, followed by RT-qPCR of the target genes CXCL8, IL1a, NFKB1A, CC20, and TNFAIP2. As expected, normalized mRNA levels were greatly reduced in RelA KD cells. In NME1 KD cells, the same genes exhibited varying degrees of reduction compared to control cells (**Figure 4D**) which agreed with RNA-seq analysis. We also assessed DNA binding by RelA using chromatin immunoprecipitation followed by qPCR (ChIP-qPCR) in scr and NME1 KD cell lines following TNFα stimulation. RelA ChIP enrichment was reduced to different extents in NME1 KD cell, suggesting these genes exhibited impaired RelA recruitment in the absence of NME1 (**Figure 4E**). Collectively, these findings suggest that NME1 is involved in regulating DNA binding and gene expression by RelA in response to TNFα.

### Expression of a subset of RelA-driven survival genes is affected in NME1-deficient unstimulated HeLa cells

The impact of NME1 on RelA on TNFα-stimulated gene regulation is not surprising given that these two proteins directly interact with each other. However, we were also interested in understanding whether NME1 plays a role in gene regulation in TNFα-independently since we identified NME1 as a facilitator of RelA’s DNA binding in unstimulated nuclear extracts. To investigate this, we examined whether reduced levels of NME1 affect the expression of RelA target genes in unstimulated conditions. Transcript levels were analyzed in control and NME1 knockdown (KD) cells under unstimulated conditions.We identified two classes of genes influenced by NME1 based on their TNFα responsiveness (**Figure 4A** and **5A**); class I genes are TNFα responsive and include the same genes discussed earlier, which are either up- or downregulated even in the absence of TNFα as shown by the volcano plot (**Figure 5B**, bottom). The class II genes are those affected by NME1-KD, either positively or negatively, and are not responsive to TNFα, and therefore may not regulated by NF-κB RelA (**Figure 5A** and **5B**, top). NME1-responsive expression of these genes in unstimulated cells suggests NME1 might act as cofactors for transcription factors other than NF-κB RelA. We were curious to examine if these genes contain κB sites in their promoters. Surprisingly, we found all of them have κB sites (**Figure 5C**). However, these κB sites are distinct from those responsive to TNFα and RelA in that the κB sites are weak and not present as clusters. As shown in **Figure 5C** and reported previously, clustered κB sites are necessary for gene regulation by RelA, and often within these clusters exist at least one κB site is a strong site (Shahabi et al., 2024).

**Figure 5.**
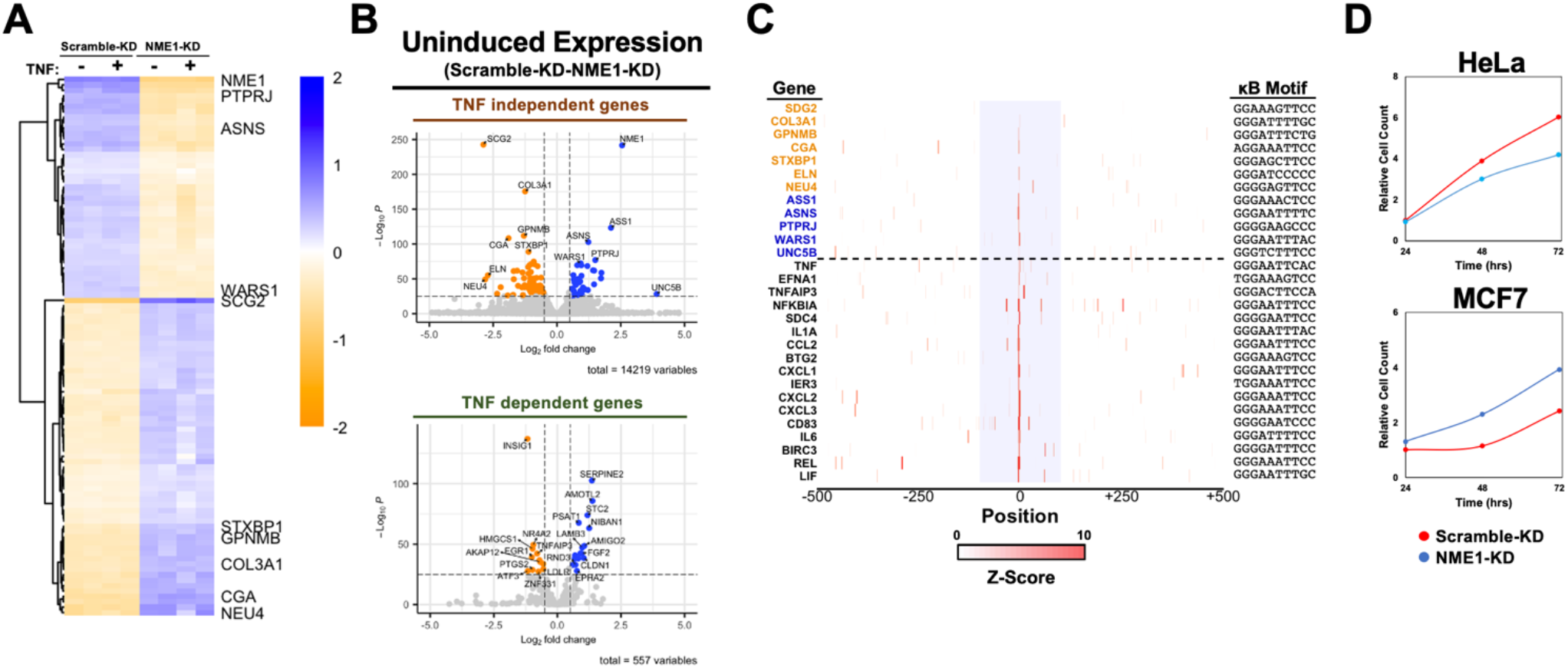
RelA-driven survival genes are regulated by NME1 in unstimulated cells. **A**. Heatmap representation of unsupervised hierarchical clustering of top 100 differentially expressed TNFα non-inducible genes from RNA-seq of scramble and NME1 knockdown HeLa cells stimulated for 1 hour (p value < 0.05). Color represents distance of log transformed transcript counts from mean. **B**. Volcano plot representation of differentially expressed TNFα-independent (top) and TNFα-dependent genes (bottom) between scramble and NME knockdown unstimulated HeLa cell lines. **C**. Identification and analysis of κB site strength and abundance at promoters of target genes. Above dotted line represents TNF-independent genes and below is TNF-dependent genes based on expression following stimulation in control cells (log_2_ fold change > 0, pvalue < 0.005). Colored genes represents increased (orange) or decreased (blue) expression following NME1-KD in uninduced cells. Central κB motif at promoter is listed on right. Highlighted blue region represents +/- 100 bp from central κB motif. **D**. Proliferation assay of HeLa (left) and MCF7 (right) in scramble and NME1 knockdown cells. Cells were seeded at 10,000 cells/well and counted every 24 hours over a total of 72 hours. Cell counts were normalized to 24 hour scrambled-KD cell count.

To assess the functional consequences of gene downregulation, we measured changes in proliferation and survival in NME1-KD-deficient HeLa cells. Cell proliferation was examined at 0, 24, 48 and 72 h post-transfection by cell count. Compared with the control KD cells, the number of viable cells in NME1 KD showed a significant decrease by 48 and 72 post cell plating (**Figure 5D**, top). Interestingly, when we tested the breast cancer cell line MCF7, we found that reduced NME1 levels unexpectedly promoted increased proliferation of these cells, suggesting that NME1 acts more in line with its established tumor suppressor role (**Figure 5D**, bottom).

## DISCUSSION

Regulation of gene expression by DNA-binding transcription factors is proving to be far more intricate than previously thought. Although mechanisms by which the DNA binding domains of these TFs recognize their DNA target sites with sequence specificity have been established through structural and mutational studies *in vitro, in vivo* regulation of this recognition process is far from being understood (Kim and Wysocka, 2023; Lambert et al., 2018; Zeitlinger, 2020). It is now clear that a transcription factor such as NF-κB as a full-length protein cannot bind its target DNA site with sufficient stability, as judged by EMSA (Mulero *et al*., 2018). NF-κB requires assistance cofactors. For instance, RPS3 and Sam68 have been clearly demonstrated to enhance DNA binding by RelA homodimer and p50:RelA heterodimer. We have recently shown that various transcription factors can also act as cofactors for RelA where they don’t require their DNA binding activity. It has also been shown that they facilitate recruitment of NF-κB to specific κB sites in cells. Both of these cofactors are also activated by stimulus and augment RelA’s DNA binding in stimulated cells (Fu et al., 2013; Wan *et al*., 2007).

In resting cells, a small amount of RelA is also present in the nucleus (Mathes et al., 2008; O’Dea et al., 2007). This constitutive presence of RelA is important for the expression of survival genes as well as other genes, such as IκBα. In this study, we show that NME1 also acts as a cofactor that supports RelA’s ability to DNA. The effect of NME1 alone on facilitating RelA’s DNA binding is modest, indicating that many other cofactors are required for full engagement which is in line with observations that RelA has many cofactors. Also supporting this notion is that each purification step removes cofactors that contribute to RelA’s DNA binding. Furthermore, our recent findings suggest that a large number of κB sites collectively recruit RelA with the help of numerous cofactors in TNFα and LPS-stimulated cells. It is plausible that different cofactors are responsible for helping RelA bind to specific subsets of κB sites. Moreover, these cofactors may interact with one another in a loosely cooperative network, allowing for combinatorial effects without sacrificing the distinct function of individual κB sites and cofactors. We also find that overlapping but distinct sets of genes require NME1 for RelA’s DNA binding in unstimulated and stimulated conditions. Further work is required to differentiate the regulatory aspects of RelA’s DNA recruitment mechanisms.

NME1 is a small nuclear protein which performs pleotropic functions, with its primary role as a dinucleotide kinase (Puts *et al*., 2018). Additionally, a related protein has been found to bind G-quadruplexes promoters (Shan et al., 2015). NME1 also stands out as the only known metazoan histidine kinase, capable of autophosphorylating His118 and transferring this phosphate to nucleotide or histidines of other proteins (Adam et al., 2020). Major interest in NME1 is due to its anti-metastatic activity. However, this activity is complex; while NME1 is anti-metastatic in cancers of epithelial origin, it can also promote the initiation in other cancers (Sharma et al., 2021; Wakefield et al., 2013; Yu et al., 2021). It will be intriguing to explore see if its transcriptional cofactor activity is tied to its anti-metastatic activity where its histidine kinase activity contributes to this dual role.

## MATERIALS and METHODS

### Antibodies and Reagents

The GST (0020), β-tubulin (0119), and IκBα (0040) antibodies were purchased from BioBharati LifeScience. The RelA (sc-372) antibody used in Western blots and ChIP assays was purchased from Santa Cruz Biotechnology. The FLAG (F1804) and control IgG ((12-371) antibodies were purchased from Sigma. The p84 (C1C3) antibody was purchased from GeneTex. The NME1 (3345) antibody was purchased from Cell Signaling. Mouse TNFα (BioBharati) was used at a final concentration of 20ng/mL for the indicated timepoints.

### Mammalian Cell Culture and Transient Transfection

HeLa and HEK293T cell lines were maintained in Dulbecco’s modified Eagle’s medium (Corning) supplemented with 10% FBS and antibiotics. For transient transfection of HEK293T, cells were plated in 6-well dishes at 80% confluence the day before transfection. The next day, 2 μg of DNA was mixed with 8 μg of polyethylenimine (PEI) in OptiMEM (Gibco) at a total reaction volume of 50 μL. After incubation at room temperature for 15 minutes, cell media was exchanged to DMEM without FBS or antibiotics and DNA:PEI complexes were added dropwise over cells and left to incubate for 4 hours at 37°C. Media was then exchanged back to DMEM supplemented with 10% FBS and antibiotics and incubated for 16 hours before harvesting. For preparation of HeLa nuclear extract, plated cells were harvested by scraping in PBS and lysed in hypototonic lysis buffer consisting of PBS supplemented with 0.1% (v/v) NP-40, 1 mM DTT, and 0.25mM PMSF. Nuclei were pelleted by centrifugation at 3000g, washed twice with ice-cold PBS, and resuspended in nuclear extraction buffer containing 25 mM Tris-HCl pH 7.5, 420 mM NaCl, 10% glycerol, 0.2 mM EDTA, 1 mM DTT, 0.5 mM PMSF, and mammalian protease inhibitor cocktail (Sigma). After incubation on ice for 30 minutes, nuclei were centrifuged at 13,000 rpm for 15 minutes at 4°C. The supernatant containing nuclear extract was collected, snap frozen on liquid nitrogen, and stored at -80°C.

The list of oligonucleotides used for generation of stable knockdown HeLa cells are listed in **Supplementary Table 1**. First, forward and reverse oligonucleotides for specific targets were annealed in buffer containing 10 mM Tris-HCl pH 7.5, 50 mM NaCl, and 1 mM EDTA by incubating in boiling water allowed to slowly cool to room temperature. Annealed DNA duplex was then ligated into the pLKO.1 TRC vector using the AgeI and EcoRI restriction sites. Lentivirus was then generated in 293T cells by cotransfecting the pLKO.1 construct with pMDLg/pRRE, pCMV-VSV-G, and pRSV-Rev expression constructs. After 48 hours, virus-containing supernatant was harvested, filtered through a 0.4 μm filter, and used to infect HeLa at a 1:10 dilution in the presence of 10 ng/μL polybrene (MilliporeSigma). After 48 hours, stably infected cells were selected by treatment of 1 μg/mL puromycin in culture media and continuously maintained in the presence of 1 μg/mL puromycin.

### Protein Expression and Purification

Expression and purification of His-tagged full-length RelA from Sf9 cells and His-tagged RelA RHR (19-304) in *E. coli* Rosetta (DE3) was performed as previously described (Li et al., 2024). His-tagged NME1 was cloned into the pET-24d vector and expressed in *E. coli* Rosetta (DE3) by growing cells to an OD600 of 0.4 and inducing with 0.25 mM IPTG overnight at room temperature. Cells were pelleted by centrifugation and lysed by sonication in lysis buffer containing 25 mM Tris-HCl pH 7.5, 300 mM NaCl, 5% glycerol, 0.1% NP-40, 10 mM imidazole, 5 mM β-mercaptoethanol, and 0.25 mM PMSF. Lysate was clarified by centrifugation at 14,000 rpm for 30 minutes at 4°C and incubated with nickel-NTA agarose beads (BioBharati LifeScience) for 2 hours at 4°C on a rotary shaker. Beads were then extensively washed with lysis buffer and His-tagged NME1 was eluted in lysis buffer without PMSF but supplemented with 250 mM imidazole. Protein quality was assessed by SDS-PAGE with Coomassie staining and peak fractions were pooled, snap frozen on liquid nitrogen, and stored at -80°C. NME1 was cloned into the pGEX-4T2 GST vector and both GST and GST-tagged NME1 were expressed in *E. coli* Rosetta (DE3). Cells were grown to an OD600 of 0.3 and induced with 0.25 mM IPTG overnight at 16°C. Cells were then pelleted by centrifugation and lysed by sonication in lysis buffer containing 50 mM Tris-HCl pH 7.5, 200 mM NaCl, 5% glycerol, 0.1% NP-40, 0.5 mM EDTA, 1 mM DTT, and 0.25 mM PMSF. The lysate was then clarified by centrifugation at 14,000 rpm for 30 minutes at 4°C and incubated with pre-equilibrated glutathione agarose beads (BioBharati LifeScience) for 2 hours at 4°C on a rotary shaker. Beads were then extensively washed with lysis buffer and eluted lysis buffer without PMSF but with 25 mM reduced glutathione (Sigma-Aldrich). Eluted protein was then dialyzed in a 6-8 kDa MWCO membrane three times in 1L of dialysis buffer containing 50 mM Tris-HCl pH 7.5, 200 mM NaCl, 5% glycerol, 0.5 mM EDTA, and 1 mM DTT and dialyzed once in 200 mL of dialysis buffer containing 50% glycerol. The dialyzed 50% glycerol stock protein was then stored at -20°C.

### Electrophoretic Mobility Shift Assay

Radiolabeled probes were incubated with the proteins under study for 20 minutes at room temperature in binding buffer containing 10 mM Tris-HCl pH 7.5, 50 mM NaCl, 10% glycerol, 1% NP-40, 1 mM EDTA, and 0.1 mg/mL poly(dI-dC). When needed, proteins were diluted in dilution buffer containing 20 mM Tris-HCl pH 7.5, 50 mM NaCl, 10% glycerol, 1 mM DTT, and 0.2 mg/mL BSA. Samples were run through a 4% nondenatured polyacrylamide gel in TGE buffer (24.8 mM Tris base, 190 mM glycine, and 1 mM EDTA) at 200V for 1 hour. Gel was then dried, exposed on a phosphor screen overnight, and scanned by Typhoon FLA 9000 imager (Cytiva).

### Fractionation and Identification of RelA-Specific Cofactors

Unstimulated HeLa nuclear extract was collected from three 15 cm dishes in a total volume of 500 μL and cleared by centrifugation at 13,000 rpm for 15 minutes at 4°C. Supernatant was collected and fractionated through a pre-equilibrated 24 mL Superose 6 (Cytiva) size-exclusion column in SEC buffer containing 25 mM Tris-HCl pH 7.5, 420 mM NaCl, 10% glycerol, 0.2 mM EDTA, 1 mM DTT, and 0.5 mM PMSF. 42 fractions at a volume of 330 μL (14 mL total volume) were collected starting 8 mL after injection, and 2 μL from fractions were tested for activity by EMSA with 5 nM of recombinant full-length RelA. Fractions 16-19 showed the most activity and were therefore combined and diluted in SEC buffer without NaCl to a final 100 mM NaCl concentration. The pooled extract was then centrifuged at 13,000 rpm for 15 minutes at 4°C and supernatant was passed through a 3 mL Mono Q (Cytiva) anion exchange column. The column was then washed with 10 mL of buffer containing 25 mM Tris-HCl pH 7.5, 100 mM NaCl, 10% glycerol, and 1 mM DTT and bound proteins were eluted with 10 mL of the same buffer but with an increasing NaCl gradient up to 700 mM NaCl. A total of 40 250 μL fractions were collected and 2 μL from fractions were tested for activity by EMSA with recombinant full-length RelA.

The fractions corresponding to Q20, Q23, Q26, Q29, and Q32 showed the highest activity and were further tested for RelA DNA-binding enhancement in an *in vitro* biotinylated DNA-pulldown assay. Biotinylated DNA was annealed as outlined previously in buffer containing 10 mM Tris-HCl pH 7.5, 50 mM NaCl, and 1 mM EDTA and immobilized onto streptavidin agarose beads (BioBharati LifeSciences) in buffer containing 25 mM Tris-HCl pH 7.5, 150 mM NaCl, 5% glycerol, 0.1% and 1 mM DTT. Beads were then washed to remove unbound DNA and mixed with 100 ng of recombinant full-length RelA at a total volume of 200 μL. 10 μL of the corresponding Mono Q fractions were added and samples were rotated for 2 hours at 4°C. Beads were then pelleted by centrifugation and washed 4 times with the same pulldown buffer. After the final wash, 4x SDS gel loading dye was added to the beads at a final dilution of 1x and beads were boiled for 10 minutes. Samples were centrifuged at 13,000 rpm for 5 minutes and supernatant was separated by SDS-PAGE and analyzed by Western blot.

Fraction Q23 showed the highest activity in both EMSA and the DNA pulldown assay and was therefore further investigated by mass-spec for peptide identification. A final pulldown was performed in buffer containing 25 mM Tris-HCl pH 7.5, 150 mM NaCl, 5% glycerol, and 1 mM DTT with 100 ng of recombinant full-length RelA, fraction Q23, and biotinylated-DNA immobilized onto streptavidin agarose beads. A control pulldown was also prepared in parallel without recombinant RelA added. The reactions were incubated overnight at 4°C with gentle rotation. Beads were then washed with the pulldown buffer 3 times and precipitated peptides were identified by liquid chromatography with mass spectrometry (LC-MS) at the UCSD Biomolecular and Proteomics Mass Spectrometry Facility.

### Luciferase Assays

Complementary oligonucleotides were first annealed by mixing at a final concentration of 2 μM in buffer containing 10 mM Tris-HCl pH 7.5, 50 mM NaCl, and 1 mM EDTA and incubating in boiling water allowed to slowly cool to room temperature. Annealed promoters were then cloned into the CMXTK-Luciferase vector (a kind gift from Dr. Chakravarti at Northwestern University Feinberg School of Medicine) at the SalI and BamHI restriction sites. HEK293T were grown in 12-well plates and transiently transfected with the FLAG-tagged overexpression construct or empty vector control, luciferase reporter DNA, and control CMV-driven Renilla. After 24 hours, cells were stimulated for 8 hours with 20 ng/mL mouse TNFα. Lysate was then collected and used for luciferase activity assay using the Dual-Luciferase Reporter Assay System (Promega). Data are represented as mean ± standard deviations (SD) of three or more independent experimental replicates.

### Pulldown Assays

NME1 was cloned into a modified pEYFP-c1 vector with YFP removed and substituted for an N-terminal FLAG tag. The day before transfection, HEK293T was plated on a 6-well plate. The next day, the cells were at approximately 80% confluence and transfected with either FLAG empty vector or FLAG-tagged NME1 with PEI. The next day, cells were washed with ice-cold PBS and lysed directly on the plate in buffer containing 25 mM Tris-HCl pH 7.5, 150 mM NaCl, 5% glycerol, 1% NP-40, 1 mM DTT, and 0.5 mM PMSF by gentle rocking at 4°C for 15 minutes. Lysate was then collected and centrifuged at 13,000 rpm for 15 minutes at 4°C. Supernatant was then collected and mixed with 50 μL of preequilibrated anti-FLAG M2 beads (Sigma) for 2 hours at 4°C on a rotary shaker. Beads were then extensively washed and mixed with 4x SDS gel loading dye to a final dilution of 1x. Samples were boiled for 5 minutes and centrifuged for 5 minutes at 13,000rpm. Supernatant was then resolved by SDS-PAGE and analyzed by Western blot.

For GST pulldown assays, first 1μg of GST or GST-tagged NME1 was mixed with 50 μL of preequilibrated glutathione agarose beads (BioBharati LifeScience) in 200 μL of pulldown buffer containing 25 mM Tris-HCl pH 7.5, 150 mM NaCl, 5% glycerol, 0.1% NP-40, and 1 mM DTT. Samples were mixed for 1 hour at 4°C with gentle rotation. Beads were then washed 4 times with pulldown buffer to wash unbound proteins and brought to a final volume of 200 μL after the last wash. 1 μg of recombinant full-length RelA and 1 μM of annealed κB or mutant κB DNA was then added and the reaction was incubated for 2 hours at 4°C with gentle rotation. Beads were then extensively washed and mixed with 4x SDS gel loading to a final dilution of 1x. Samples were then boiled for 5 minutes and centrifuged at 13,000rpm for 5 minutes. Supernatant was then resolved by SDS-PAGE and analyzed by Western blot.

### RNA Isolation and Real-Time qPCR

A list of oligonucleotides used in RT-qPCR reactions is listed in **Supplementary Table 1**. HeLa was plated in 6 well dishes and total RNA was isolated the next day with TRIzol (Invitrogen) and purified by isopropanol precipitation following the manufacturers recommendations. RNA concentration was determined by nanodrop and cDNA was synthesized in a 5 μL reaction from 500 ng of RNA using SuperScript IV VILO Master Mix (ThermoFisher). The cDNA was then diluted 1:4 to a total volume of 20 μL and 1 μL was used as template for qPCR with the Luna qPCR Master Mix (New England Biolabs) in a total reaction volume of 10 μL. Values were normalized to GAPDH and data are represented as mean ± standard deviation of three independent experimental replicates.

### Chromatin Immunoprecipitation qPCR

A list of oligonucleotides used in ChIP-qPCR experiments is listed in **Supplementary Table 1**. For every two immunoprecipitation reactions, a confluent 10 cm dish of HeLa was used. Cells were first treated with TNFα or DMSO for 30 minutes. Formaldehyde was then added directly to the media at a final concentration of 1% and incubated on the cells for 10 minutes at room temperature with gentle rocking. Crosslinking was then quenched by addition of 125 mM glycine and incubation at room temperature for 5 minutes with gentle rocking. Cells were then washed twice with ice-cold PBS and collected by scraping in 1mL of PBS. Cells were then pelleted by centrifugation and resuspended in 1 mL of PBS supplemented with 0.1% NP-40, 1 mM DTT, and 0.25 mM PMSF. The cell pellet was then gently pipetted 5 times to facilitate cytoplasmic lysis and nuclear fractionation, and nuclei was then pelleted by centrifugation. The nuclear pellet was then resuspended in 1 mL of RIPA buffer containing 50 mM Tris-HCl pH 7.5, 150 mM NaCl, 2 mM EDTA, 1% NP-40, 0.1% sodium deoxycholate, 0.1% SDS, 0.5 mM PMSF, and mammalian protease inhibitor cocktail (Sigma). Resuspended nuclei were sonicated on ice with a micro tip sonicator (Branson) to generate DNA fragments with an average length of 500 bp. Sonicated nuclear extract was then centrifuged at 13,000 rpm for 15 minutes at 4°C and supernatant was precleared with 25 μL protein AG PLUS agarose beads (BioBharati LifeScience) and 500 ng of IgG control antibody (ThermoFisher) for 1 hour at 4°C with rotation. Extract was then centrifuged at 13,000 rpm for 10 minutes at 4°C, and precleared supernatant was divided into two equal reactions and mixed with 25 μL of protein AG PLUS agarose beads and 500 ng of either IgG or anti-RelA antibody. Immunoprecipitation reactions were incubated at 4°C overnight with gentle rotation. Beads were then washed three times for 5 minutes each with 1 mL of RIPA buffer, then RIPA buffer supplemented with 500 mM NaCl, RIPA buffer with NaCl substituted with 250 mM LiCl, and lastly with TE buffer containing 10 mM Tris-HCl pH 7.5 and 1 mM EDTA. Immune complexes were then eluted from beads in 150 μL of elution buffer containing 1% SDS and 100 mM sodium bicarbonate pH 8.0 for 30 minutes at room temperature with rotation. Eluted complex was then mixed to 6 μL of 5 M NaCl and 2 μL of 100 mg/mL RNase A (Qiagen) and incubated overnight at 65°C to reverse cross-linking and digest RNA. The next morning, 2 μL of Proteinase K (Invitrogen) was added and incubated at 60°C for 1 hour. DNA was then extracted with phenol:chloroform:isoamyl alcohol (25:24:1) (Invitrogen) and isopropanol precipitation following manufacturers recommendation. DNA pellet was resuspended in 50 μL of TE buffer and 1 μL was used in qPCR reaction with Luna qPCR Master Mix (New England Biolabs).

### RNA Sequencing and Analysis

Scramble and NME1 knockdown HeLa cell lines were initially plated on 6 well plates. The next day, duplicate wells from each cell line were treated for 1 hour with either DMSO or 20 ng/mL TNFα. Cells were then washed once with ice-cold PBS and RNA was isolated with TRIzol (Invitrogen) following manufacturers recommendations. RNA quality was then assessed by TapeStation (Agilent) and RNA with a RIN score greater than 8.0 was further processed. Poly-A enriched libraries were prepared from 1 μg of total RNA using the mRNA HyperPrep Kit (KAPA) with unique dual-indexed adapters (KAPA) following manufacturers recommendations. Library quality was assessed by DNA TapeStation (Agilent) and quantified with Qubit 2.0 fluorometer (Life Technologies). Libraries were pooled and underwent paired-end sequencing using the NovaSeq6000 (Illumina) at the UCSD Institute for Genomic Medicine (IGM). For analysis of sequencing data, read quality was first checked by FASTQC. Reads were then mapped to human genome using OSA/Oshell (Omicsoft). Reads were then normalized and differentially expressed genes were analyzed using DESeq2 (v1.38.3).

## ACKNOWLEDGEMENTS

The work was supported by NIH grant GM085490 to GG. SS was supported by the CMG training grant during his PhD. Authors thank the Ghosh lab members for helpful discussion.

## AUTHOR CONTRIBUTIONS

G.G. conceived the original idea. S.Shahabi further refined the idea and conducted most experiments. M.M. performed RNA-seq analysis. S.S. supervised bioinformatic section of the manuscript. G.G. wrote the original draft which was modified by S.Shahabi. All other authors read the manuscript.

## FIGURES and FIGURE LEGENDS

**Supplemental Figure 1.**
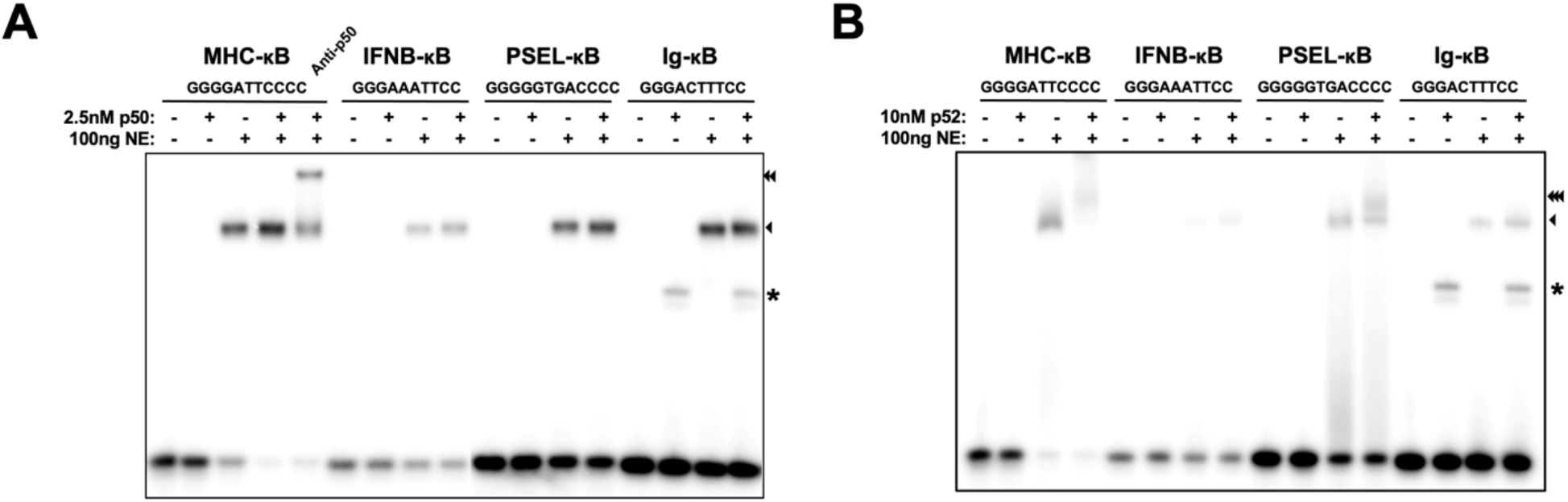
**A**. and **B**. EMSA assay with recombinant p50 (**A**.) or p52 (**B**.) homodimers and unstimulated HeLa nuclear extract. DNA sequences used in reaction are indicated above (Asterisk is non-specific binding from nuclear extract, single arrow is binding of p50 or p52 homodimers to κB DNA, double arrow is supershifted band with p50-specific antibody, and triple arrow is probable p52:Bcl3 complex).

**Supplemental Figure 2.**
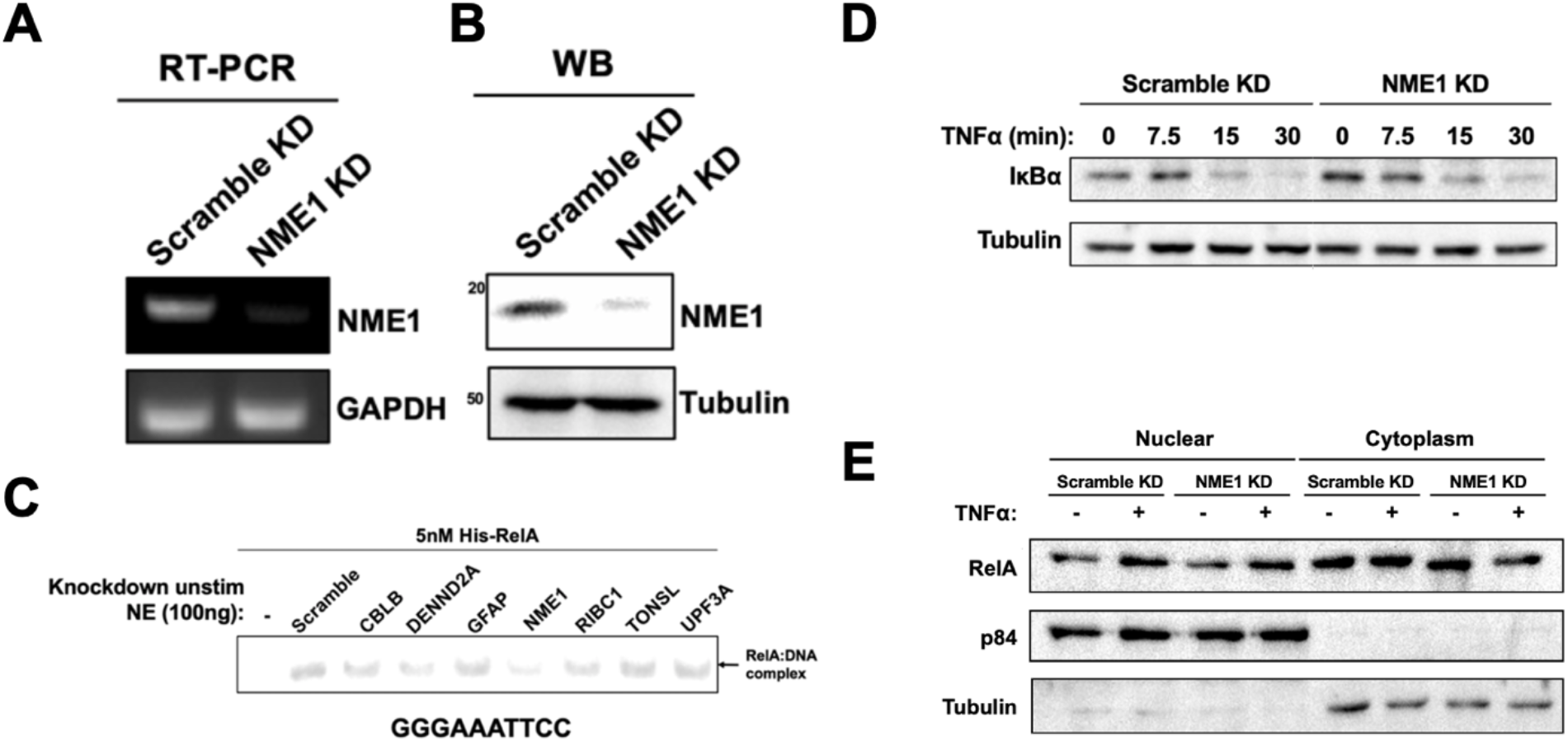
**A**. and **B**. RT-PCR (**A**.) and Western blot (**B**.) analysis of scramble and NME1 knockdown stable HeLa cell lines. **C**. EMSA analysis with 5 nM recombinant FL-RelA and 100 ng of indicated unstimulated nuclear extract from stable knockdown HeLa cell lines. The κB DNA sequence used in EMSA is indicated below. **D**. Stable scramble and NME1 knockdown HeLa cell lines were stimulated with TNFα for the indicated timepoints and IκBα degradation was analyzed by Western blot of whole cell lysate. **E**. Stable scramble and NME1 knockdown HeLa cell lines were stimulated with TNFα for 15 minutes. Cells were then fractionated and nuclear and cytoplasmic fractions were analyzed by Western blot using the indicated antibodies.

**Supplemental Figure 3.**
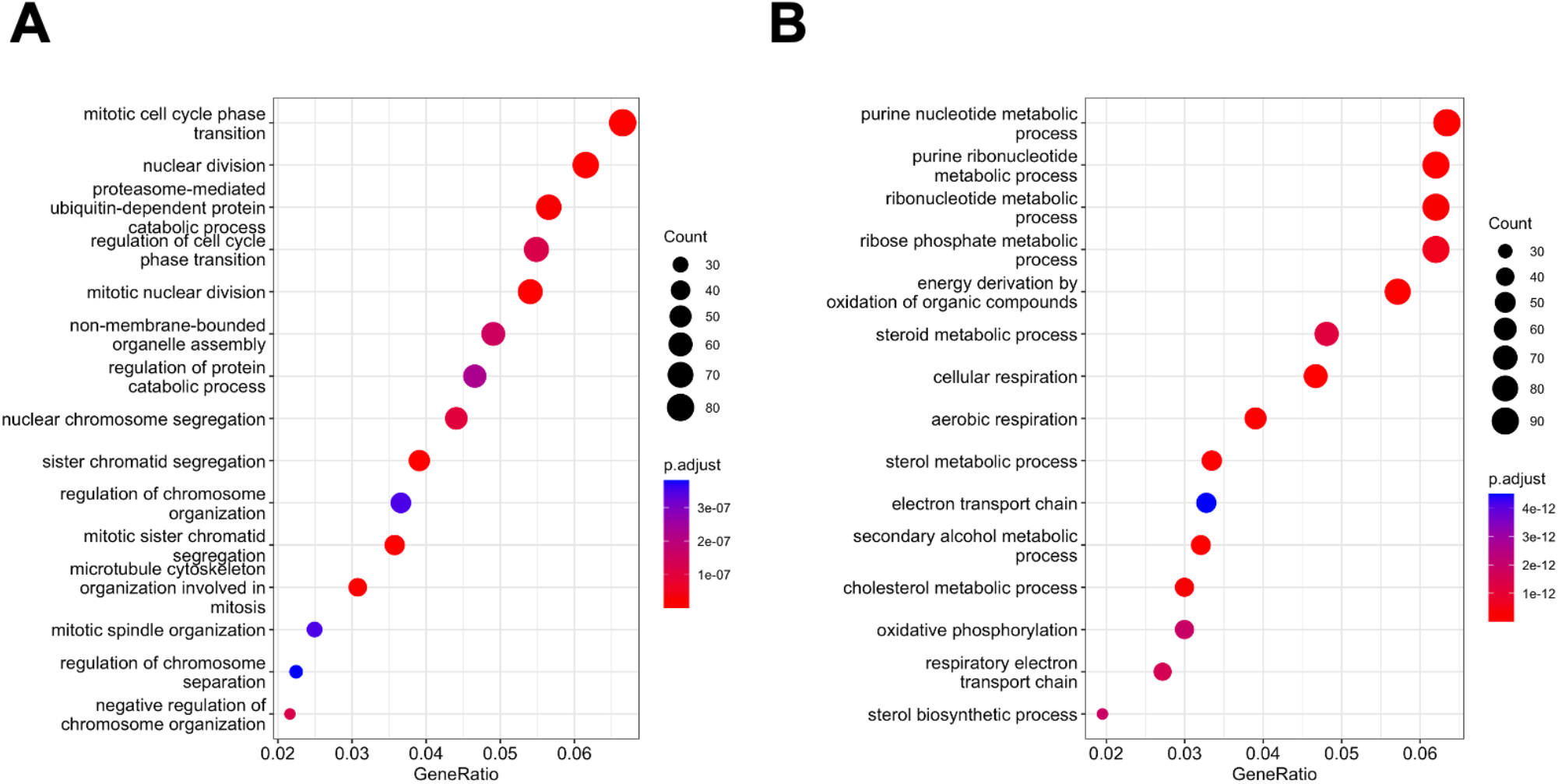
**A**. and **B**. Gene ontology terms enriched with genes downregulated (**A**.) or upregulated (**B**.) upon NME1 knockdown in unstimulated cells identified using clusterProfiler.

## REFERENCES

Adam, K., Ning, J., Reina, J., and Hunter, T. (2020). NME/NM23/NDPK and Histidine Phosphorylation. Int J Mol Sci 21. 10.3390/ijms21165848.

Fu, K., Sun, X., Zheng, W., Wier, E.M., Hodgson, A., Tran, D.Q., Richard, S., and Wan, F. (2013). Sam68 modulates the promoter specificity of NF-kappaB and mediates expression of CD25 in activated T cells. Nature communications 4, 1909. 10.1038/ncomms2916.

Ghosh, G., Wang, V.Y., Huang, D.B., and Fusco, A. (2012). NF-kappaB regulation: lessons from structures. Immunological reviews 246, 36–58. 10.1111/j.1600-065X.2012.01097.x.

Ghosh, S., and Hayden, M.S. (2012). Celebrating 25 years of NF-kappaB research. Immunological reviews 246, 5–13. 10.1111/j.1600-065X.2012.01111.x.

Hoffmann, A., Levchenko, A., Scott, M.L., and Baltimore, D. (2002). The IkappaB-NF-kappaB signaling module: temporal control and selective gene activation. Science (New York, N.Y.) 298, 1241–1245. 10.1126/science.1071914.

Jin, X., Ding, D., Yan, Y., Li, H., Wang, B., Ma, L., Ye, Z., Ma, T., Wu, Q., Rodrigues, D.N., et al. (2019). Phosphorylated RB Promotes Cancer Immunity by Inhibiting NF-kappaB Activation and PD-L1 Expression. Molecular cell 73, 22-35.e26. 10.1016/j.molcel.2018.10.034.

Kim, S., and Wysocka, J. (2023). Deciphering the multi-scale, quantitative cis-regulatory code. Molecular cell 83, 373–392. 10.1016/j.molcel.2022.12.032.

Lambert, S.A., Jolma, A., Campitelli, L.F., Das, P.K., Yin, Y., Albu, M., Chen, X., Taipale, J., Hughes, T.R., and Weirauch, M.T. (2018). The Human Transcription Factors. Cell 172, 650–665. 10.1016/j.cell.2018.01.029.

Leung, T.H., Hoffmann, A., and Baltimore, D. (2004). One nucleotide in a kappaB site can determine cofactor specificity for NF-kappaB dimers. Cell 118, 453–464. 10.1016/j.cell.2004.08.007.

Li, T., Shahabi, S., Biswas, T., Tsodikov, O.V., Pan, W., Huang, D.B., Wang, V.Y., Wang, Y., and Ghosh, G. (2024). Transient interactions modulate the affinity of NF-kappaB transcription factors for DNA. Proceedings of the National Academy of Sciences of the United States of America 121, e2405555121. 10.1073/pnas.2405555121.

Lin, J., Kato, M., Nagata, K., and Okuwaki, M. (2017). Efficient DNA binding of NF-kappaB requires the chaperone-like function of NPM1. Nucleic acids research 45, 3707–3723. 10.1093/nar/gkw1285.

Mathes, E., O’Dea, E.L., Hoffmann, A., and Ghosh, G. (2008). NF-kappaB dictates the degradation pathway of IkappaBalpha. The EMBO journal 27, 1357–1367. 10.1038/emboj.2008.73.

Mulero, M.C., Shahabi, S., Ko, M.S., Schiffer, J.M., Huang, D.B., Wang, V.Y., Amaro, R.E., Huxford, T., and Ghosh, G. (2018). Protein Cofactors Are Essential for High-Affinity DNA Binding by the Nuclear Factor kappaB RelA Subunit. Biochemistry 57, 2943–2957. 10.1021/acs.biochem.8b00158.

Mulero, M.C., Wang, V.Y., Huxford, T., and Ghosh, G. (2019). Genome reading by the NF-kappaB transcription factors. Nucleic acids research. 10.1093/nar/gkz739.

O’Dea, E.L., Barken, D., Peralta, R.Q., Tran, K.T., Werner, S.L., Kearns, J.D., Levchenko, A., and Hoffmann, A. (2007). A homeostatic model of IkappaB metabolism to control constitutive NF-kappaB activity. Mol Syst Biol 3, 111. 10.1038/msb4100148.

Pan, W., Meshcheryakov, V.A., Li, T., Wang, Y., Ghosh, G., and Wang, V.Y. (2023). Structures of NF-kappaB p52 homodimer-DNA complexes rationalize binding mechanisms and transcription activation. Elife 12. 10.7554/eLife.86258.

Puts, G.S., Leonard, M.K., Pamidimukkala, N.V., Snyder, D.E., and Kaetzel, D.M. (2018). Nuclear functions of NME proteins. Lab Invest 98, 211–218. 10.1038/labinvest.2017.109.

Shahabi, S., Biswas, T., Shen, Y., Zhou, Y., Sanahmadi, R., and Ghosh, G. (2024). Cooperativity among clustered kB sites within promoters and enahncers dictaes transcriptional specificity of NF-kB RelA along with specific cofactors. bioRxiv. 10.1101/2024.07.03.601930.

Shan, C., Lin, J., Hou, J.Q., Liu, H.Y., Chen, S.B., Chen, A.C., Ou, T.M., Tan, J.H., Li, D., Gu, L.Q., and Huang, Z.S. (2015). Chemical intervention of the NM23-H2 transcriptional programme on c-MYC via a novel small molecule. Nucleic acids research 43, 6677–6691. 10.1093/nar/gkv641.

Sharma, S., Sengupta, A., and Chowdhury, S. (2021). Emerging Molecular Connections between NM23 Proteins, Telomeres and Telomere-Associated Factors: Implications in Cancer Metastasis and Ageing. Int J Mol Sci 22. 10.3390/ijms22073457.

Siggers, T., Chang, A.B., Teixeira, A., Wong, D., Williams, K.J., Ahmed, B., Ragoussis, J., Udalova, I.A., Smale, S.T., and Bulyk, M.L. (2011). Principles of dimer-specific gene regulation revealed by a comprehensive characterization of NF-kappaB family DNA binding. Nature immunology 13, 95–102. 10.1038/ni.2151.

Todeschini, A.L., Georges, A., and Veitia, R.A. (2014). Transcription factors: specific DNA binding and specific gene regulation. Trends Genet 30, 211–219. 10.1016/j.tig.2014.04.002.

Wakefield, A., Soukupova, J., Montagne, A., Ranger, J., French, R., Muller, W.J., and Clarkson, R.W. (2013). Bcl3 selectively promotes metastasis of ERBB2-driven mammary tumors. Cancer Res 73, 745–755. 10.1158/0008-5472.Can-12-1321.

Wan, F., Anderson, D.E., Barnitz, R.A., Snow, A., Bidere, N., Zheng, L., Hegde, V., Lam, L.T., Staudt, L.M., Levens, D., et al. (2007). Ribosomal protein S3: a KH domain subunit in NF-kappaB complexes that mediates selective gene regulation. Cell 131, 927–939. 10.1016/j.cell.2007.10.009.

Wang, V.Y., Huang, W., Asagiri, M., Spann, N., Hoffmann, A., Glass, C., and Ghosh, G. (2012). The transcriptional specificity of NF-kappaB dimers is coded within the kappaB DNA response elements. Cell Rep 2, 824–839. 10.1016/j.celrep.2012.08.042.

Yu, L., Wang, X., Zhang, W., Khan, E., Lin, C., and Guo, C. (2021). The multiple regulation of metastasis suppressor NM23-H1 in cancer. Life Sci 268, 118995. 10.1016/j.lfs.2020.118995.

Zeitlinger, J. (2020). Seven myths of how transcription factors read the cis-regulatory code. Curr Opin Syst Biol 23, 22–31. 10.1016/j.coisb.2020.08.002.

